# Superstitious learning of abstract order from random reinforcement

**DOI:** 10.1101/2022.02.02.478909

**Authors:** Yuhao Jin, Greg Jensen, Jacqueline Gottlieb, Vincent P. Ferrera

**Affiliations:** Dept. of Biological Sciences, Columbia University; Dept. of Psychology, Reed College; Dept. of Neuroscience, Columbia University; Kavli Institute for Brain Science, Columbia University; Zuckerman Mind Brain Behavior Institute, Columbia University

**Keywords:** learnability detection, spurious learning, transitive inference, reinforcement learning

## Abstract

Survival depends on identifying learnable features of the environment that predict reward, and avoiding others that are random and unlearnable. However, humans and other animals often infer spurious associations among unrelated events, raising the question of how well they can distinguish learnable patterns from unlearnable events. Here, we tasked monkeys with discovering the serial order of two pictorial sets: a “learnable” set in which the stimuli were implicitly ordered and monkeys were rewarded for choosing the higher-rank stimulus and an “unlearnable” set in which stimuli were unordered and feedback was random regardless of the choice. We replicated prior results that monkeys reliably learned the implicit order of the learnable set. Surprisingly, the monkeys behaved as though some ordering also existed in the unlearnable set, showing consistent choice preference that transferred to novel untrained pairs in this set, even under a preference-discouraging reward schedule that gave rewards more frequently to the stimulus that was selected less often. In simulations, a model-free RL algorithm (*Q*-learning) displayed a degree of consistent ordering among the unlearnable set but, unlike the monkeys, failed to do so under the preference, discouraging reward schedule. Our results suggest that monkeys infer abstract structures from objectively random events using heuristics that extend beyond stimulus-outcome conditional learning to more cognitive model-based learning mechanisms.

---

Learning is vital for survival. Learning mechanisms have been extensively studied in laboratory tasks that are learnable i.e., contain objective regularities that can be discovered through trial and error. However, natural environments are complex spaces that expose learners both to learnable tasks and random associations. For example, upon seeing a red Toyota driving by in the rain one could, in principle, wonder if there is a relationship between red Toyotas and rain. Because learning entails significant costs (in terms of effort, time, risk and missed opportunities), animals would ideally distinguish learnable tasks – to which they may wish to devote their resources and interest – from random associations that are “unlearnable” – which they should ignore [1].

It is unclear, however, how well animals can make this distinction. Converging evidence shows that humans [2, 3, 4, 5, 6, 7, 8] and non-human animals [9, 10, 11, 12] learn spurious associations in variety of conditions. Spurious learning is generally explained in terms of simple associative learning that overestimates causal relationships between external events or between the animal’s actions and outcomes [13, 14, 15, 16]. Indeed, simple associative learning models, like the Rescorla–Wagner model, depend heavily on correlations between events and can be easily fooled into strengthening associations based on incidental coincidences [17]. However, since animals also infer more elaborate structures, an open question is if they also inappropriately impose complex structures on objectively random events.

Here, we examined this question in the context of a “transitive inference” (TI) task in which monkeys were required to infer the ordinal relationships among a set of stimuli that had a hidden order (e.g., “ABCDE”). Subjects received pairs of stimuli from that ordered set and were rewarded for choosing the higher rank stimulus. By training subjects on adjacent pairs (e.g., AB, BC, etc.) and testing them on non-adjacent pairs (e.g., AC, BD, etc.) we examined if they obeyed the law of transitivity by rapidly generalizing the inferred order to untrained pairs. The task is well-suited to our question because it has been extensively characterized in multiple species [18, 19, 20, 21] and conclusively shown to require mechanisms beyond reward associations [22, 23, 24]. In our current task, monkeys received pairs from an ordered, learnable set as in the classical TI task and, in randomly interleaved trials, pairs from an unlearnable image set that had no hidden order and the subjects’ choices were rewarded randomly. We analyzed if choices on each stimulus set were consistent with a hidden order, how they were affected by different reward schedules and if they could be reproduced by associative *Q*-learning algorithms.

## RESULTS

### Subjects learned latent order by trial and error

In each trial, rhesus monkeys (*n* = 3) saw two pictures and touched one to proceed. The pictures that were available for each trial were both drawn from one of two sets of 5 pictorial stimuli. The stimuli in one set had an underlying rank-ordering and subjects were rewarded for choosing the stimulus that had a higher rank in each pair (**Fig. 1A, top**, the letter denotes the rank with A the highest and E the lowest rank). The stimuli in the other set were unordered and subjects were rewarded probabilistically regardless of their choices (**Fig. 1A, bottom**, the letter only represents serial number of each stimulus but not rank). Because the objectively ‘correct’ order of the stimuli could be learned from trial-and-error feedback for the former but not the latter set, we refer to the sets as, respectively, “learnable” (L) and “unlearnable” (U).

**Figure 1.**
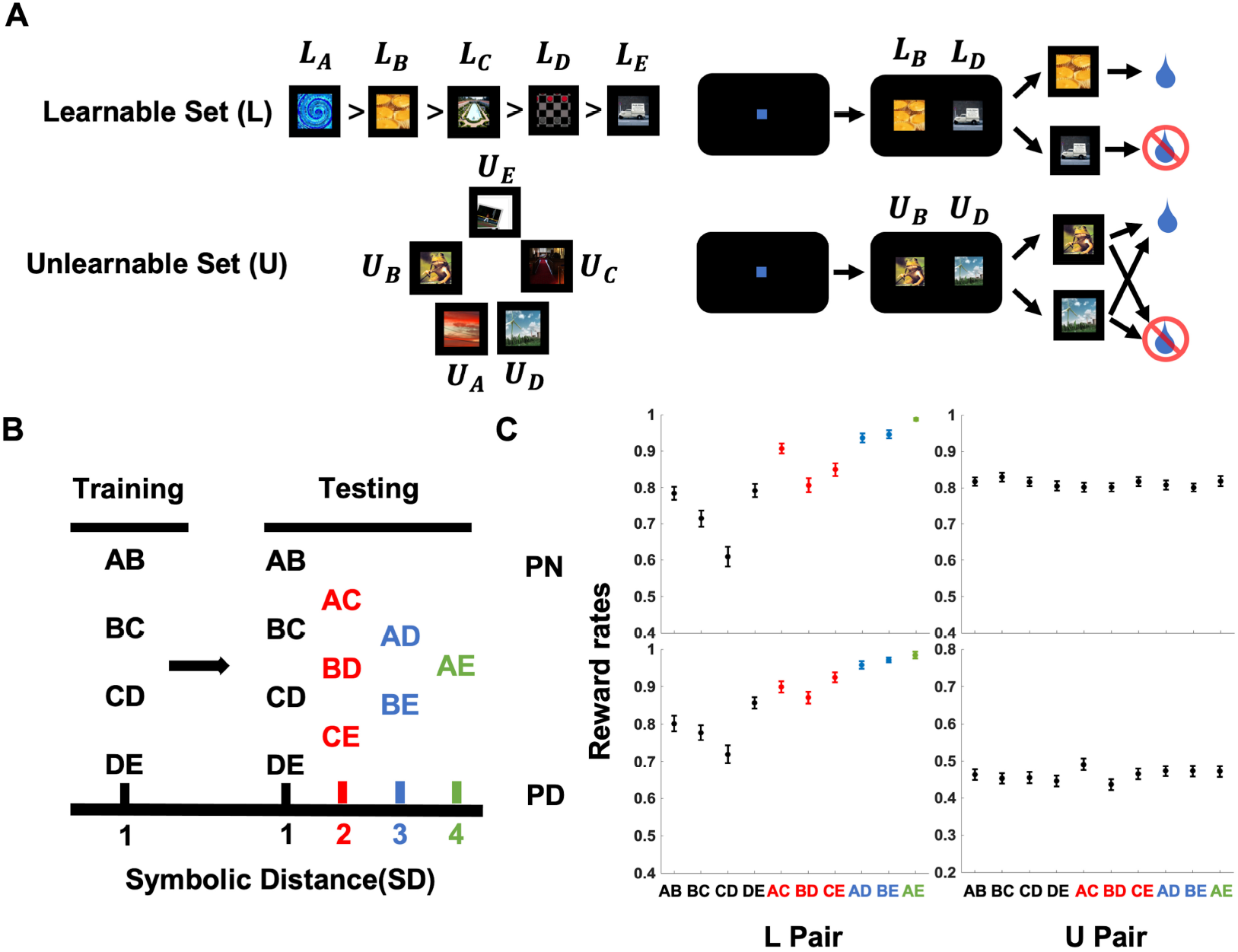
Task paradigm. **A.** Subjects were tasked with discovering the implicit ordering of sets comprising 5 pictorial stimuli. In “learnable” sets (L), the stimuli were assigned an order that could be inferred by trial and error. In learnable trials, the picture-outcome association was consistent and predictable: the picture with a higher rank was always associated with reward, whereas the other was not (top); In “unlearnable” sets (U), there was no pre-defined order and feedback was delivered probabilistically. In unlearnable trial, either response could potentially result in reward. Under the preference neutral (PN) condition, reward probabilities for U pairs were yoked to recent performance on L pairs; Under the preference discouraging (PD) condition, the reward probability for each stimulus was inversely related with how recently it had been selected. See Methods for details. **B.** Subjects were first presented with training blocks consisting of only the adjacent pairs (SD=1). After training, they transitioned to testing blocks consisting of all the possible pairs (SD=1 to 4). The transition from training to testing let us evaluate performance on novel pairs that rely on transitive inferences. **C.** Reward rates over all the L and U pairs across subjects. Error bars denote the standard error of the means.

During each session, the subjects interacted with a new set of L and U stimuli different from other sessions. L and U trials were randomly interleaved and there was no explicit cue signaling which set the stimulus pair was drawn from. The sessions consisted of a training phase, during which subjects only experienced the four adjacent stimulus pairs from the L set and 4 randomly selected pairs from the U set, followed by a testing phase with all possible pairs (**Fig. 1B**). This design allowed us to examine if the subjects inferred an order during training and spontaneously transferred it to the new pairs during testing.

To better understand how subjects approached the U trials, each session used one of two different reward schedules. Under the “preference neutral” schedule (PN), the reward probability for U pairs was equated to that for L pairs by dynamically adjusting it to match the mean reward rate for L pairs on the preceding 10 L trials (while remaining independent of which U stimulus was chosen). Under the second, “preference discouraging” reward schedule (PD), the reward probability for each U stimulus was inversely related to how recently the subject had selected it. The PD schedule thus discouraged repeated choice of any specific U stimulus, and yielded maximal rewards if differences in U stimuli preferences were minimized and each U stimulus was selected equally (see Methods for details). Consistent with the reward schedule design, the reward rates for the U and L sets were indistinguishable for the PN schedule (L: .7581 ± .0107, U: .7473 ± .0109), but differed appreciably for the PD schedule (L: .7801 ± .0121, U: .4626 ± .0035) (see also **Fig. S1A**).

As expected from previous studies, all the subjects reliably learned the order of the L sets, shown by above-chance response accuracies and by a robust symbolic distance effect (SDE; reward rates increases as the difference in rank between the two stimuli presented on each trial grows) that was significant on average (**Fig. 1C left**, PN: F(3, 236) = 93.391, *p* < .001; PD: *F*(3, 236) = 76.34, *p* < .001) and in each individual monkey (**Fig. S1B**); In contrast, for the U set, average reward rates were equal for all pairs in both reward schedules (**Fig. 1C right**, PN: *F*(3, 236) = 0.747, *p* = 0.524; PD: *F*(3, 236) = 0.917, *p* = 0.432, SD applied here is calculated as the difference in serial number between the two stimuli in each U pair, **Fig. S1B**).

### Subjects chose as if imposing an order on unordered stimuli

Despite the lack of objective order among the U stimuli, subjects seemed to treat them as if they were ordered. **Fig. 2A (top)** shows this result for an example session with the PN reward schedule, in which the subject developed a clear choice preference in the L set (which was consistent with the objective stimulus label) and also a clear preference within the U set (despite the arbitrary stimulus labels). Under the PD schedule, a similar result for the L set was seen and U set still manifested a certain degree of differential preferences over stimuli (**Fig. 2A, bottom**).

**Figure 2.**
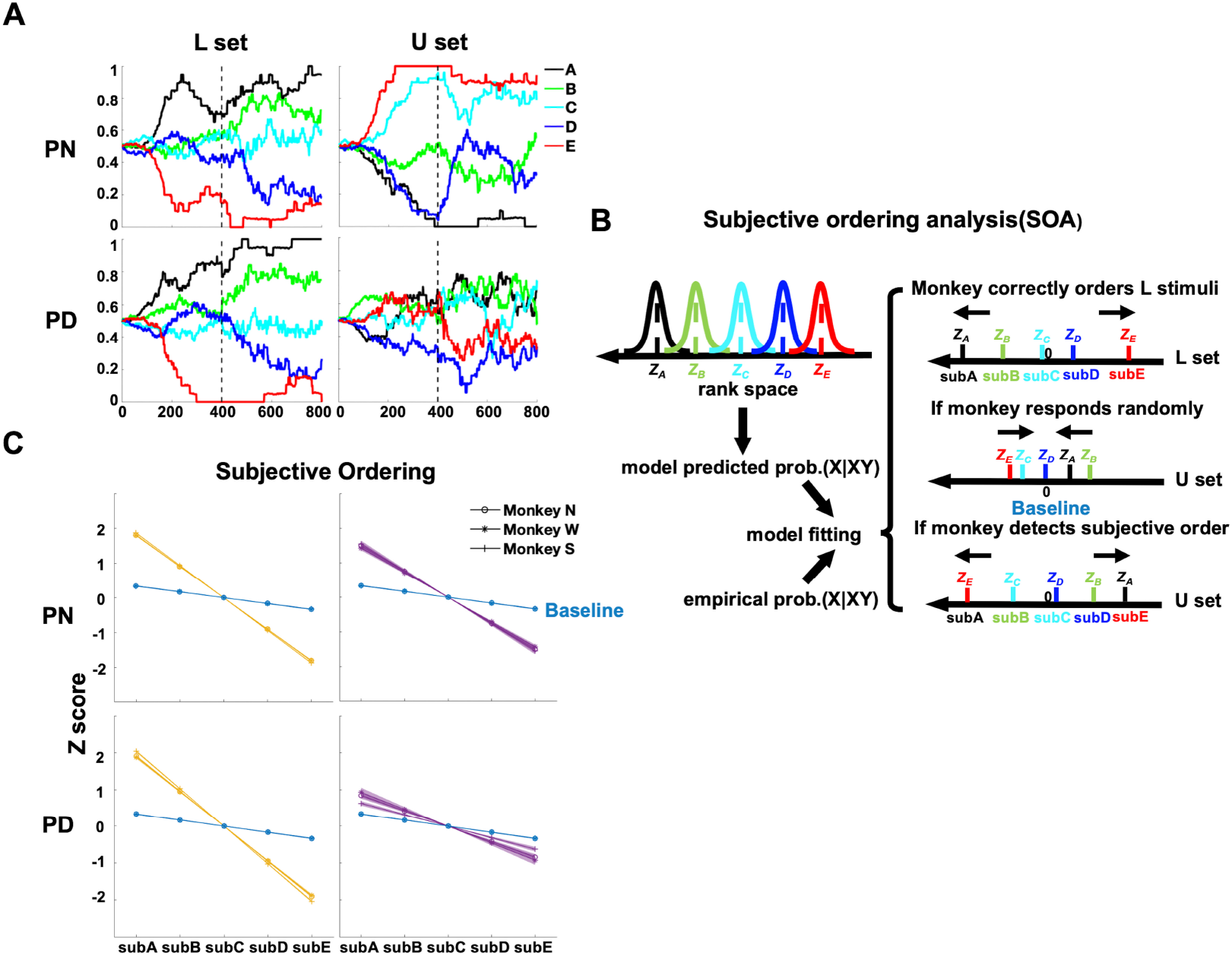
Subject created abstract ordering upon the unlearnable stimuli. **A.** Example session showing moving averages across trials (*n*=80 for training and *n*=50 for testing) of choice frequency for over L and U stimuli. **B.** A schematic depiction of the subjective ordering analysis (SOA). The model assumed that a subject represented each stimulus as a position along a linear continuum with normally distributed uncertainty. Each stimulus had its own mean (*z*-score) and SD parameter. The model was used to infer the most probable z-scores for each stimulus given the subject’s history of choices across all pairs. All stimuli could be subjectively labelled subA to subE based on their z-scores. The L set was expected to display a subjective order that was consistent with the true order (top). If a subject selected U set items randomly, the inferred *z*-scores would shrink together close to zero (middle). However, if subject created a subjective order, then the *z*-scores would spread out to reduce their overlapping uncertainties (bottom). Baseline estimates for the z-scores were obtained by simulating random responding, in order to have a rigorous null against which to compare behavior. **C.** Estimated subjective item positions for both L and U sets under both reward schedules during the testing phase. All subjects showed stronger preference orderings (i.e. steeper slopes) than baseline. Error bars represented the standard errors of the means.

Even subjects choose randomly, the probability of choosing each stimulus may not be perfectly at chance level, which will incur some degree of preferred ordering. To test this, and to estimate the strength of these apparent preferences among U stimuli, we used a model-based subjective ordering analysis (SOA) that fit choices in each session based on the assumption that subjects represented stimuli along some continuum with some uncertainty about stimulus positions (**Fig. 2B**; Methods; [22]). For data acquired during the testing phase, the analysis produced a *z*-score indicating the relative rank of each stimulus. A stronger *z*-score gradient indicated stronger preference, more consistent choices and less overlap between inferred stimulus ranks. For the L-sets, the gradients (slopes) over the *z*-scores were significantly higher than would be expected from a baseline of random responding during the testing phase (PN: all *p* < .001; PD: all *p* < .001, rank-sum test, **Fig. 2C left**). Furthermore, the gradients for the L sets were equivalent whether stimuli were ranked according to the objective ordering defined by the experiments or according to the subjective ordering estimated by the analysis (testing phase: PN: all *p* > .2; PD: all *p* > .7, rank-sum test). Put another way, when an objective ordering existed, the SOA analysis of behavior reliably recovered the true stimulus ranks from each subject’s preferences, confirming the validity of the SOA analysis.

The *z*-score gradients for U sets for all 3 subjects displayed slopes that were significantly steeper than baseline in both PN and PD schedule (**Fig. 2C right**, PN: all *p* < .001; PD: all *p* < .001, rank-sum test). This suggests that subjects displayed consistent preferences among U stimuli, despite rewards that were either independent of stimulus (in the PN schedule) or actively discouraged preferences (in the PD schedule). However, slopes were lower in PD schedule than in the PN schedule (all *p* < .001, rank-sum test). Evidence for subjective ordering among U stimuli was also found during training (**Fig. S2A**). The strength of the subjective ordering did not systematically change across sessions (**Fig. S2B**) and was not correlated with the gradient in L sets (**Fig. S2C**), suggesting these preferences are stable over the long term and unrelated to engagement with the task. Thus, the monkeys’ preferences appeared to reveal a tendency to impose order on unordered stimulus sets, even under a reward schedule in which such preference incurred a cost by reducing the rate of reward.

### Subjects transferred the subjective ordering from training to testing

Comparisons of training and testing stages for the U sets showed that the subjective preferences that developed during training remained consistent during the testing stage. **Fig. 3A** illustrates this result for a representative subject under PN and PD schedules, in which the rank order preference for stimuli during testing (shown by the color labels) was the same as the rank ordering during training (shown by the relative position of the traces). Two analyses verified this result quantitatively. First, the *z*-scores over each U stimulus estimated from training and testing data were significantly correlated for the PN schedule (**Fig. S3A**, *r* > .59, *p* < .001) and for the PD schedule (**Fig. S3A**, *r* > .06, *p* < .01). Second, during testing, subjects showed a robust subjective symbolic distance effect (sub SDE, distances were coded based on the subjective ordering inferred during training) for U sets that was similar to the objective symbolic distance effect (obj SDE, distances were coded based on objective order) for the L sets (**Fig. S3B**, **Fig. 3B**). This further confirms that they were choosing as if there were a consistent order. A logistic regression revealed slopes that were significantly higher relative to the subSD randomized null case for each subject in the PN schedule (**Fig. 3B, top right**, U 95% CI for the subSD regressor N: [0.3675, 0.3707], W: [0.4065, 0.4100], S: [0.4564, 0.4604]; subSD regressor shuffled N: [0.0010, 0.0040], W: [0.0421, 0.0452], S: [−0.0216, −0.0182]) and for two of three subjects in the PD schedule (**Fig. 3B**, bottom right, U 95% CI for the subSD regressor N: [0.1588, 0.1618], W: [0.1298, 0.1328], S: [0.0154, 0.0183]; shuffled N: [−0.0437, −0.0408], W: [0.0298, 0.0327], S: [−0.0372, −0.0343]). Thus, subjective orderings estimated using training data could be used to make predictions about performance for novel U pairs presented during the testing phase.

**Figure 3.**
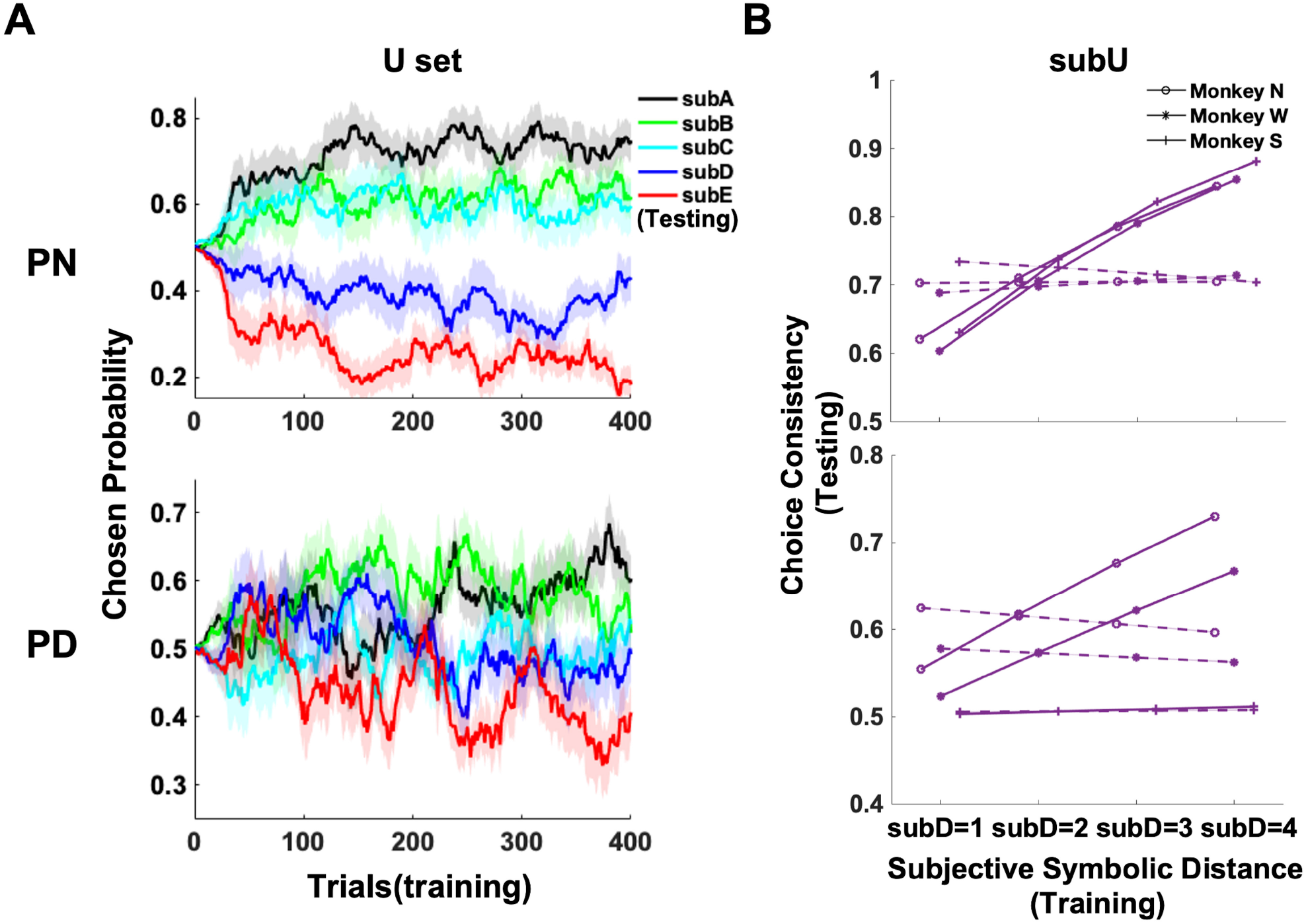
Subject transferred the subjective order on the U set from training to testing. **A.** Moving averages across trials (n=16) of choice frequency of each U stimulus during training for an example subject. Here, the subA-subE labels were determined using the testing data. Shaded regions correspond to the bootstrapped 95% CI. **B.** Logistic regression estimates of preference as a function of symbolic distance between the subjective ranks of U stimuli at the start of testing. Solid lines denote subject estimates, whereas dashed lines represent a null case with the distance shuffled. Error bars denote the standard errors over all the posterior Bayesian regression coefficients. However, the error bars are invisible because it’s smaller than the size of the lines.

### *Q*-learning does not explain observed behavior

A difficulty in evaluating behavior under an ostensibly “uninformative” condition is that subjects may draw spurious conclusions from the random feedback being provided. Put another way, reinforcement learning (RL) models can sometimes form preferences even when feedback is random and uninformative. To evaluate this possibility, we simulated behavior using the model-free *Q*-learning RL algorithm [25], which is often considered a canonical example of reward prediction error learning. In addition to having a ‘learning rate’ parameter, our *Q*-learning implementation used a softmax decision rule [26, 27] with a ‘temperature’ parameter that governed how much random variation was introduced into decisions. Posterior parameter distributions for each subject were estimated numerically using the Stan programming language [28, 29], as described in the Methods. As expected, *Q*-learning succeed in learning the veridical order in L set and showed matching reward rates between L and U sets under PN schedule but decoupled reward rates under PD schedule (**Fig. S4A**).

While *Q*-learning developed preferences (and thus a subjective ordering) among U stimuli when rewarded using the PN schedule, it failed to display reliable preferences under the PD schedule. Under the PN schedule, applying the SOA analysis to the model choices showed that *Q*-learning produced significant subjective ordering for the U sets relative to baseline of random response, in a manner similar to the monkeys (**Fig. 4A**, averaged behavior: top, mean slope Baseline: −0.1671; *Q*-learning: −0.7314; 95% CI Monkey: [-0.7503,-0.7375], **Fig. S4B top**). Applying the transfer analysis showed that *Q*-learning produced significant transfer from training to testing, despite weaker than that shown by the monkeys (**Fig. 4B** averaged behavior: top, U subSD regressor 95% CI Monkey: [0.4095, 0.4143], mean *Q*-learning: 0.2234; subSD regressor shuffled 95% CI Monkey: [0.0068, 0.0107], mean *Q*-learning: 0.0092, **Fig. S4C top**). However, under the PD schedule *Q*-learning performed worse than baseline in forming subjective orderings (**Fig. 4A**, averaged behavior: bottom, mean slope Baseline: −0.1671; *Q*-learning: −0.1211; 95% CI Monkey: [-0.4058,-0.3916], **Fig. S4B bottom**) and showed no transfer effect (**Fig. 4B**, averaged behavior: bottom, U subSD regressor 95% CI Monkey: [0.0997, 0.1059], mean *Q*-learning: −0.0026; subSD regressor shuffled 95% CI Monkey: [-0.0176, −0.0135], mean *Q*-learning: 0.0028, **Fig. S4C bottom**). Therefore, our model simulation result showed that the subjective ordering extends beyond standard model-free reinforcement learning algorithms (*Q*-learning) and the mechanism is needed to be investigated in the future.

**Figure 4.**
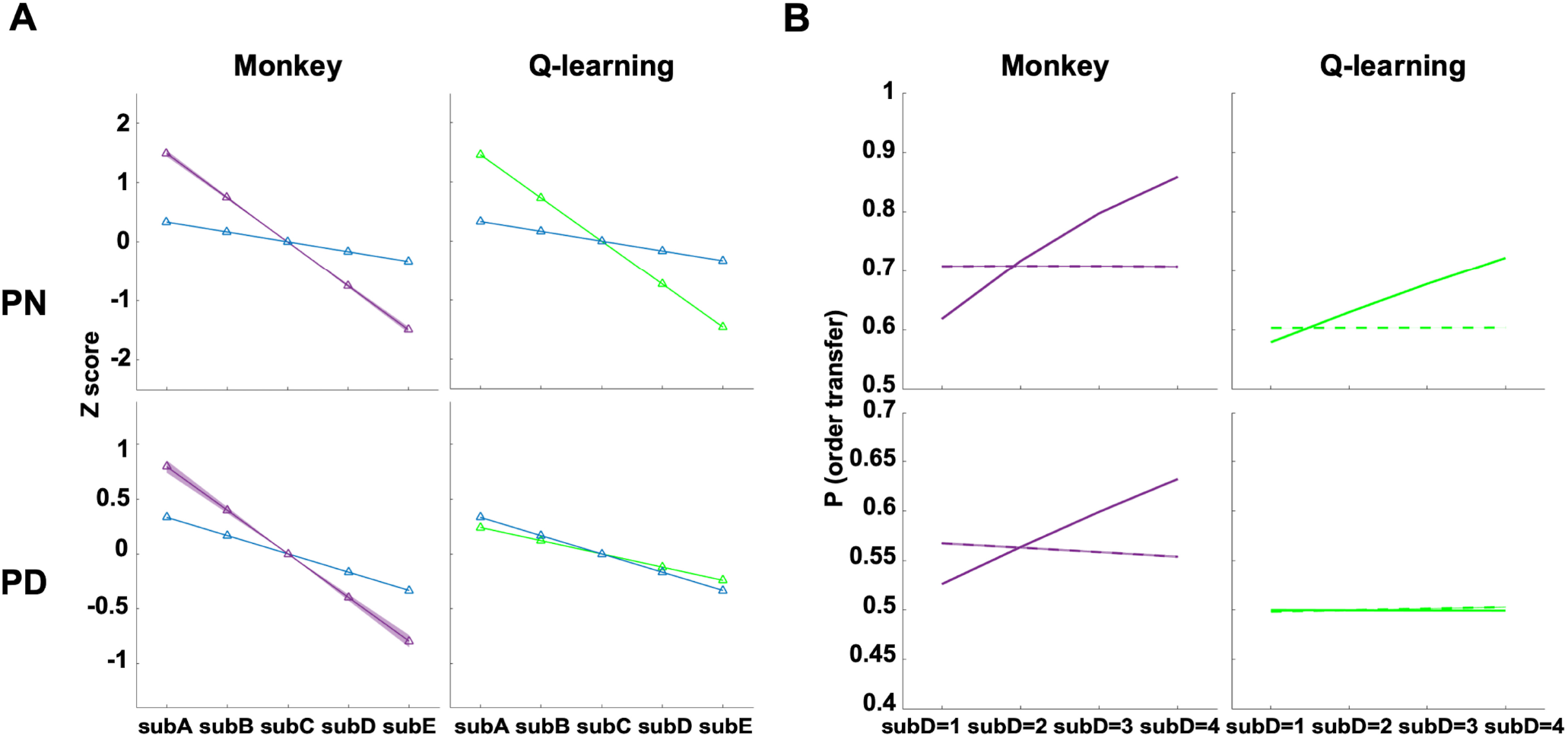
*Q*-learning cannot explain monkey’s behavior. **A.** Subjective ordering analysis of RL simulations using *Q*-learning, as compared with that of subjects. **B.** Mean performance at the start of testing, sorted by symbolic distance (solid lines) and a null case (dashed lines). Error bars denote the standard errors over all the posterior Bayesian regression coefficients.

## DISCUSSION

While the mechanisms underlying learning of structured tasks have been intensively explored, little is known about how animals react to unstructured, random events. Here we provide evidence that animals do not always effectively distinguish between learnable and unlearnable tasks. We exposed non-human primates to unordered sets of stimuli within the context of a TI paradigm, and discovered that they treated the sets as though they were ordered. Subjects developed preferences consistent with the stimuli being rank ordered, and these “subjective orderings” persisted throughout the session and were predictive of choices made to newly introduced pairs.

Importantly, we show that subjective orderings were inferred through mechanisms that go beyond simple associative learning. We show that, for unlearnable sets, in stark contrast with learnable sets, subjects were insensitive to past feedback and showed equal probability of win-stay and lose-stay strategies (**Fig. S5**, [5, 6, 11]). Moreover, choices consistent with subjective ordering remained strong in a preference discouraging (PD) schedule that actively discouraged them by dynamically increasing reward probability for whichever alternative had been selected least often. In contrast, *Q*-learning algorithm failed to produce subjective ordering under this schedule, although it replicated it under a preference-neutral schedule that did not discourage consistent preferences, capturing the well-known vulnerability of associative models to spurious reward correlations [13, 14, 15, 16, 17]. These results are consistent with a wealth of studies showing that pure associative learning is not sufficient to explain TI learning [22, 23, 24, 30]. Together, we show that monkeys ascribe subjective structure to objectively random events based on inferential mechanisms that are in part model-based or at least, rely on more complex assumptions than retrospective reward maximization.

The mechanisms generating subjective ordering are unknown, but we hypothesize they may be of two kinds. One possible mechanism involves generalization. Extrapolating from their experience with serial learning and other learnable sets, subjects may have *a priori* assumed that all the sets they experience in a session have a learnable order. Thus, if subjects begin the task believing that each stimulus is pre-assigned to a certain rank, it stands to reason that this *a priori* representation would be resilient against at least some counterfactual information. This view is consistent with previous evidence that when monkeys and humans were shown the “derived” pairings consisting of each stimulus from distinct trained ordered set, they relied on the known ranks held by each stimulus in their original set to judge the novel combination, suggesting that they spontaneously assume that the two sets use the same ranking scale even when there is no logical necessity for this being the case [31, 32, 33]. More broadly, this view is consistent with the proposed role of generalization heuristics in guiding exploration in complex contexts under high uncertainty [34, 35, 36].

A second, not mutually exclusive mechanism, is related to uncertainty and ambiguity, which may pose cognitive costs. Humans and monkeys are generally aversive to uncertainty, ambiguity and conflict [37, 38, 39] and in humans, choices among food items with similarly high value are associated with higher anxiety relative to choices among more distinct items [40]. This suggests that making decisions under higher uncertainty has affective and cognitive costs and attempts to avoid these costs by reinterpreting choice situations may result in irrational behaviors such as causal illusions and superstitious choices [41, 42]. Thus, the monkeys’ assumption that unlearnable sets were ordered may have been motivated by a desire to reduce subjective uncertainty of making the choice despite the external feedback being stochastic. The relative roles of generalization and attitudes to uncertainty will be important areas for future investigations.

Whatever the eventual mechanism turns out to be, our result that subjective ordering persisted through the PD schedule when it reduced reward rates suggests that it is a powerful phenomenon that may lead to suboptimal choices. This may be particularly important in “strategic learning” scenarios, in which learners must decide how to allocate time and effort among competing learning activities. A recent report showed that humans devoted disproportionate effort to random and unlearnable tasks at the expense of improving on alternative learnable tasks [8]. A theoretically efficient method for avoiding “randomness traps” is to value competing activities in proportion to learning progress – the extent to which one’s success rates improve over time [1, 43]. However, this strategy may be considerably weakened if people, similar to the monkeys we studied, were less sensitive to actual reward rates – or, more precisely, to the contingency between their choices and outcomes – and acted more based on their assumption that a structure exists. Furthermore, our study sheds light on the interplay between the strength of subjective ordering and the changes of reward rates. Subject showed significantly stronger subjective ordering and transfer effect for the U set in the PN schedule than in the PD schedule. Under the PN schedule, U set feedback was yoked to that of L set and thus the reward rate tended to grow as subjects learned the L set ordering. Under the PD schedule, the feedback always yielded lower reward rates for any stimuli subjects began to prefer and thus reinforce uniform selection among the stimuli (**Fig. 2C**, **Fig. 3B**). However, while subjective orderings were weaker under the PD schedule, they were still reliably detectable. Future studies may thus probe the importance of two factors as we proposed – a *priori* beliefs about the existence of learnable structure and experienced changes in reward rates – in judging learnability and allocating time among competing learning activities. Future work could also try this approach on humans and even allow them to freely choose between learnable and unlearnable trial to test whether humans, as our results hint, creates persistent superstitious structure from random feedback.

## MATERIALS AND METHODS

### Subjects

Subjects were three adult male rhesus macaques (*Macaca mulatta*), N, W and S. All subjects had different amounts of prior experience with the classic transitive inference task (single ordered set, transfer paradigm) from years (N and S) to only a month (W). However, none of the subjects had been exposed to the unordered U sets, or to the PN and PD schedules. Subjects were water restricted for maintaining high motivation. Subjects earned the reward by receiving water drops with each drop having a volume of about 0.1ml. Typical performance per session yielded between 150ml and 300ml. Subjects were also provided a ration of biscuits each day before the task and fruit as extra bonus after the task. The study was implemented obeying to the guidelines provided by the Guide for the Care and Use of Laboratory Animals of the National Institutes of Health (NIH). This work was also approved by the Institutional Animal Care and Use Committees (IACUCs) at Columbia University.

### Apparatus

Subjects performed the task by using touchscreen connected to a computer (Windows 10) while sitting on the chair. The touchscreen (Elo Touch Solutions, California) presented subjects with a 15” by 12” HD display (1280×1024 resolution at 60Hz), both showed the objects and recorded the response. Tasks were programmed in Matlab (2018a, Mathworks, Natick MA) using Psychophysics Toolbox [44]. To deliver rewards, the computer sent out the commend to the Arduino Uno interface, which then relayed the signal to the solenoid valve with 0.1ml being output through a tube installed on the primate chair per time the valve was open.

### Procedure

Pictorial stimuli were selected at random from a large bank of stock photographs and further processed to equalize their size (250 × 250 pixels). Sets were examined in advanced to confirm that they did not include stimuli that could be easily confused for one another. Each day, subjects performed one session and presented with two pictorial sets with 5 stimuli each. Different sets were used over days and across subjects. Over the two sets, one set is learnable (L), meaning the stimuli were pre-assigned an arbitrary rank order and subject was required to learn to infer the veridical order by trial and error (denoted as *L_A_, L_B_, L_C_, L_D_, L_E_*). Another set is unlearnable (U), meaning the stimuli were not ordered and subject would acquire zero knowledge of order because of the random feedback (denoted as *U_A_, U_B_, U_C_, U_D_, U_E_*, **Fig 1A**).

During each trial, two stimuli (pair) were shown side-by-side on the touchscreen. Both stimuli were drawn either from the L set or the U set. Subject should touch one of the stimuli to complete the trial. At the beginning of each trial, a solid blue square (100 × 100 pixels) was presented at the center of the screen to gain the subject’s attention and guide their hands to touch the square to initiate the trial within 3 seconds after the square appeared, otherwise the current trial would be aborted and the same trial would be repeated again until the response was made. After the initiation, the blue square disappeared and the randomly drawn stimulus pair was shown at an equal distance (289 pixels) from the center. Subject had four seconds to make a decision by touching one of the stimuli; otherwise the current trial would be skipped and missed forever and the task would move on to the next trial. If the pair came from the L set, subject would receive positive feedback (check sign, followed by 2 drops of reward and sound cue) or negative feedback (cross sign, followed by 3 seconds time-out with the screen being dark) based on whether the decision accorded with the order in the L set. Therefore, choosing the stimulus with higher rank will always result in positive outcome. If the current stimulus pair came from the U set, the feedback would be delivered probabilistically independent of which pair of stimuli were displayed.

Each subject was exposed to two schedules that kept the design for L set the same but varied for U set with the same number of sessions (20 for each subject each schedule). Under the Preference Neutral (PN) schedule, the reward probability for each U trial was the same between two shown stimuli and yoked to the average reward rates over the ten past L trials to the current trial. Under the Preference Discouraging (PD) schedule, the reward probability over all stimuli started from 0.3 when each was presented the first time, but was inversely correlated with the recent chosen frequency over all the subsequent presentations by following the equation (1 – (1 – 0.3)^(*n*+1)^) where *n* denotes the number of trials that the stimulus remained unchosen since the last time it was chosen. Therefore, the degree of preference over all U stimuli did not affect the reward rates under PN schedule since it is totally dependent on the subject’s performance on the L trials whereas PD schedule punished persistent preference over any U stimulus because any repetitive selection of the same stimulus would result in lower reward rates than intermittent selection indicative of choosing each stimulus evenly.

Each session comprised multiple adjacent pairs blocks (training, N and S: 25; W: 50), followed by multiple all pairs blocks (testing, all subjects: 10). Each training block contained 16 trials (4 different stimulus pairs per set (AB, BC, CD, DE) × 2 stimulus sets × 2 for spatial counterbalancing) and each testing block contained 40 trials (10 different stimulus pairs from each set × 2 stimulus sets × 2 for spatial counterbalancing) (**Fig 1B**). Within each block, the sequence of the trials was randomized with L and U trials interleaved to ensure monkey could not predict the future trials. Overall, N and S completed up to 800 trials per session, whereas W completed up to 1200 trials.

### Data Analysis

#### Subjective Ordering Analysis (SOA)

To best infer the subjective ordering over all L/U stimuli and evaluate the ordering strength, we assumed a model-based representation where each stimulus *i* normally distributed along the same positional continuum with its own mean *μ_i_* and variance 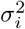. When a given pair (*XY*) was presented, the stimulus with higher mean was more likely to be chosen. For example, the probability to choose *X* was given by:

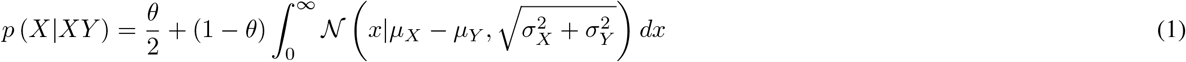

Here, *θ* denotes the degree to which subject ignored the current presentation and made random responses. Thus, subjects had probabilities that could be computed simultaneously for four pairs (during training) or ten pairs (during testing) per session in each schedule (we only considered *p* (*A|AB*) but not *p* (*B|AB*) since *p* (*A|AB*) = 1 – *p* (*B|AB*) and so on for other pairs). These simultaneous equations were solved using Bayesian multilevel model fitting using Markov chain Monte Carlo (MCMC) method in Stan programming language [28, 29]. For more details on this procedure, see [22].

After model fitting, each parameter *μ_i_* was *z*-scored relative to other position estimates; that it, the mean value of all stimuli in a set was centered at zero. The subjective ordering was manifested by performing linear regression over the sorted z-scores in descending order, by which we could use the slope to measure the ordering strength and sorted *z*-scores to label the subA-subE over each set of L/U stimuli. Since even random response are still expected to display a non-zero slope due to the sorting step, we simulated sessions in which each subject responded entirely randomly. Then each session of the simulated data went through the aforementioned subjective ordering procedure, which gave us the “baseline” slopes that were expected from thus null model of random responding, in order to better evaluate how far that the subjective ordering is from random responding.

#### Logistic Regression

To look at whether subject carried the subjective ordering from training to testing stage, we applied the following logistic regression model:

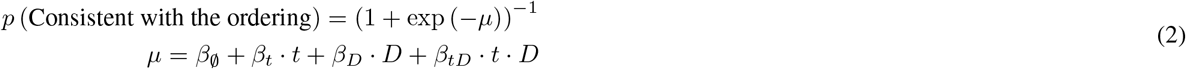

In the regression model, the response variable represented whether subject’s or artificial agent’s choice during testing accorded to the veridical order in L set or the subjective order in U set during training (Boolean output). Such dependent variable indicating ordering transfer was predicted by trial number (*t*), symbolic distance (*D*, based on ground truth order for the L set and subjective ordering for the U set) and their interaction, yielding three slope terms *β_t_, β_D_*, and *β_tD_*. The intercept term 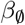 and *β_D_* served to estimate *p* (Consistent with the ordering) at trial zero with respect to different *D* for both sets. Therefore, typical symbolic distance effect would show a significant positive *β_D_* .In other words, a larger value of *β_D_* corresponds to a preference that is more consistent to the order from training, and thus that the subjective order transferred from training to testing. Both t and D were centered at zero to decrease the slope covariances before Bayesian multi-level model fitting performed in Stan programming language using MCMC method [28, 29]. Additionally, a separate analysis was performed in which *D* was shuffled over trials, sessions, and subjects for both L and U set respectively, in order to provide a control case and help evaluate the significance of the SDE.

#### *Q*-Learning Simulation

We used model-free reinforcement learning algorithm *Q*-learning to examine the hypothesis that the subjective ordering could be solely driven by evaluation of each stimulus’s reward value under the random feedback. Since each trial was totally independent of each other, the agent’s action *a_t_* on trial *t* had no effect on the next state *S*_*t*+1_. Therefore, the *Q*-learning model implied for both sets were identical to the Rescorla-Wagner model [45]:

Value Updating Rule:

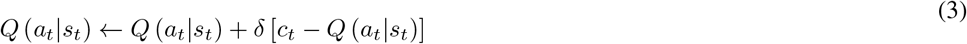

Action Policy:

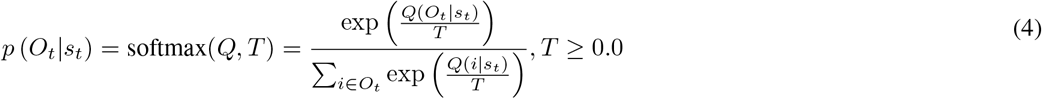

In the model, *s_t_* was the stimulus pair in each trial, *a_t_* was the agent’s choice, *c_t_* was the actual reward that agent received, the probability that subject chose certain option *O_t_* was calculated by the softmax function. The *Q*-learning model implemented here followed the asymmetrical updating rule assuming that only the *Q* value of the chosen stimulus got updated. Three hyperparameters: *δ*(learning rate, determines how fast the value gets updated), *T* (temperature, decides how much variability to introduce into choices) and *γ* (spatial bias, +1 for left response and −1 for right response) underwent Bayesian multi-level model fitting performed in Stan programming language using MCMC method [28, 29]. The mean values of the best fitting parameters were shown as followed: PN: L set *δ*: N 0.0097 W 0.0036 S 0.0155; T: N 0.0788 W 0.0527 S 0.0671; *γ*: N 0.3713 W −0.0879 S 0.4620; U set *δ*: N 0.0152 W 0.0082 S 0.0280; T: N 0.0726 W 0.0639 S 0.0785; *γ*: N 0.5957 W −0.1008 S 1.1292; PD: L set *δ*: N 0.0125 W 0.0040 S 0.0108; T: N 0.0849 W 0.0445 S 0.0413; *γ*: N 0.1437 W −0.6379 S 0.7093; U set *δ*: N 0.4601 W 0.5285 S 0.3048; T: N 15.8807 W 22.1249 S 0.9469; *γ*: N 0.5722 W −0.6415 S 2.3307. The parameters indicated that for the U set, temperature were much higher under PD (*T* close to or higher than 1) than PN schedule (*T* smaller than 1), indicating that subjects’ decisions relied less on the retrospective reward history because of the high volatility of the feedback under PD schedule. Next, we simulated the *Q*-learning behaviors with 1500 sessions per subject under each schedule, which were thereby used for SOA analysis and logistic regression. It was noteworthy that the response variable no longer referred to single trial (binary outcome) but rather to a percentage of trials consistent to the order out of a collection of trials pooled over all sessions that shared the same regressor value (*t, D*, or both). Such binomial regression would largely lower the computational cost.

#### Past-trial effect analysis

To evaluate how subject’s current decision was adaptive to the past-trial feedback, we quantified the percentage of win-stay and lose-stay for each L and U pair separately from the testing stage session by session for each subject and schedule, calculated as the chance of staying on the stimulus in pair XY that was rewarded/punished when last time XY was presented. Lastly, we averaged the percent of win-stay and lose-stay over all L and U pairs respectively.

## Acknowledgments

We would like to thank Dr. Allain-Thibeault Ferhat and Yvonne Li for their feedback on drafts of this manuscripts.

## Author Contributions

The study was designed and the manuscript was prepared by YJ, GJ, JG & VPF. Data were collected and analyzed by YJ.

## SUPPLEMENTAL FIGURES

**Figure S1.**
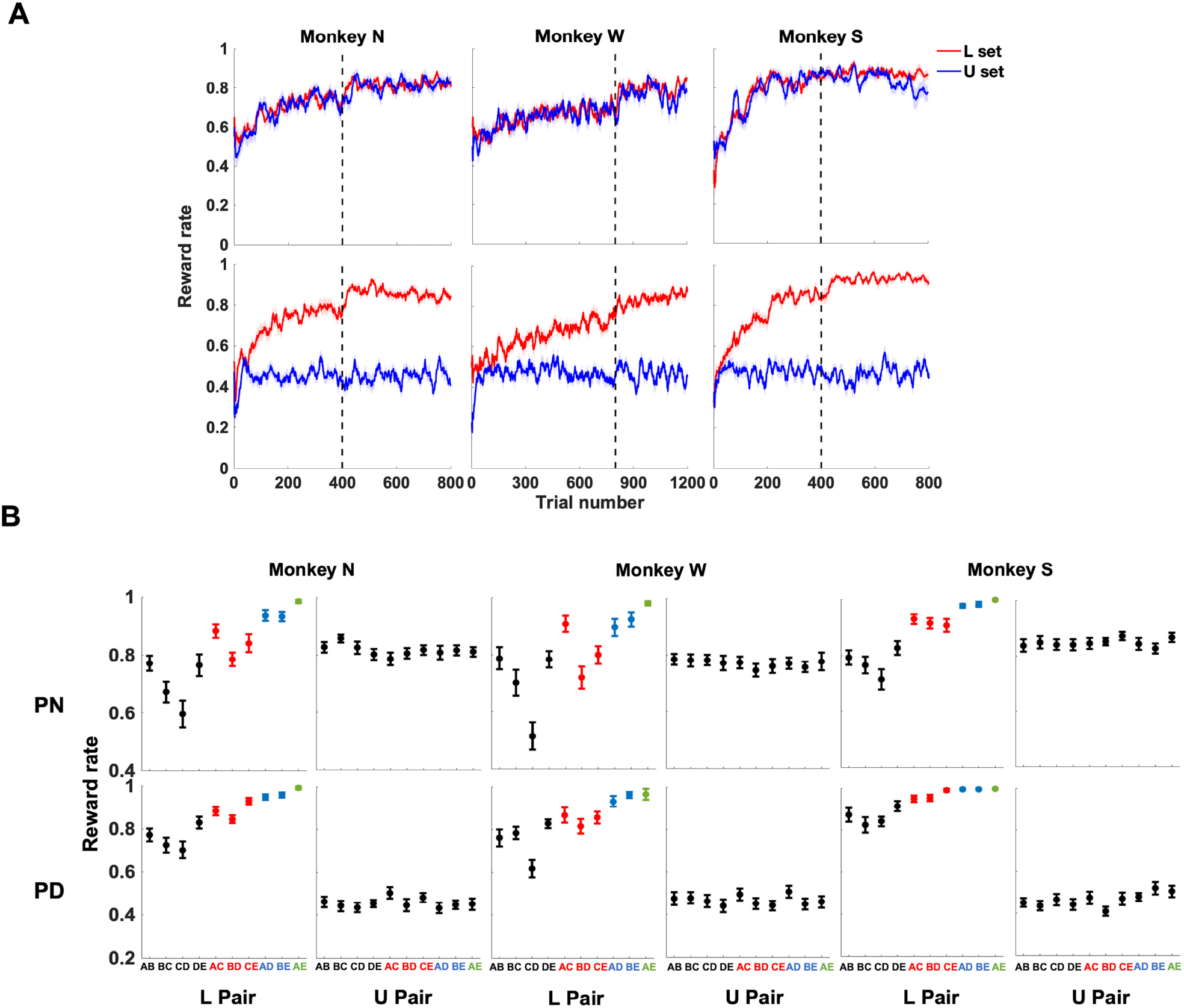
Monkeys consistently correctly inferred the order of L sets. **A.** The learning curve calculates 10 L/U trials rolling reward rates over the course of the session. It showed that all three monkeys’ performances on the L set started from chance to certain asymptotic high level under both schedules because each session subject encountered different stimulus set and had to learn from the scratch. The dash line demarcates the transition from training to testing stage. **B.** Reward rates over all the L and U pairs in individual subject (PN: L set, all *F*(4, 76) > 22, *p* < .001; U set, all *F*(4, 76) < 1.02, *p* > .38; PD: L set, all *F*(4, 76) > 23, *p* < .001; U set, all *F*(4, 76) < 2.8, *p* > .04).

**Figure S2.**
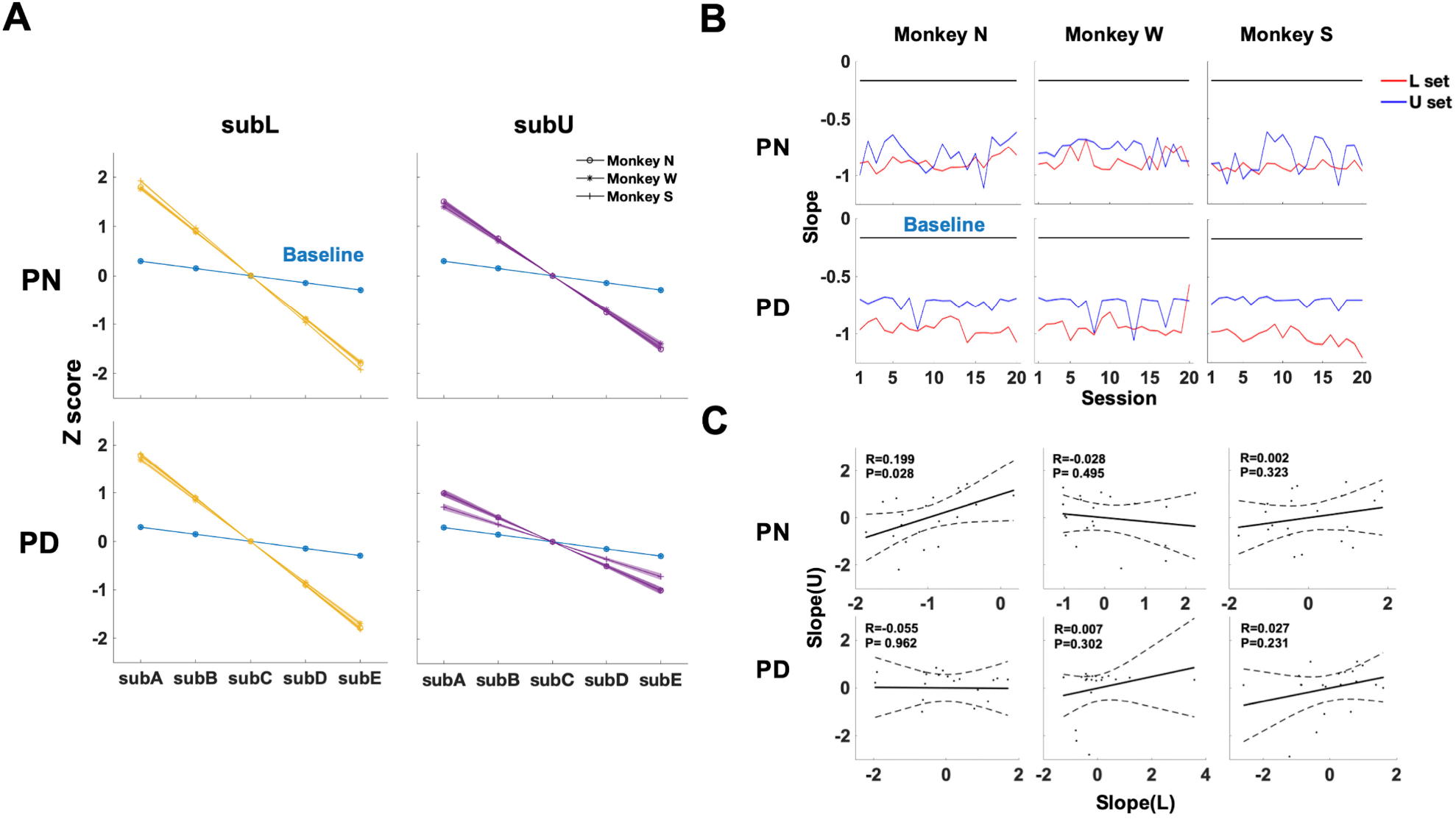
The subjective ordering is decorrelated with the amount of sessions trained and the task engagement. **A.** Estimated subjective item positions for both L and U sets under both reward schedules during the training phase. Error bars denote the standard errors of the means (PN: L set all *p* < .001, U set all *p* < .001; PD: L set all *p* < .001, U set all *p* < .001, PN vs PD: U set all *p* < .001, rank-sum test). **B.** Plotting the slopes from L and U stimuli separately over sessions for each subject, which reflects session-wise change of the ordering strength. Error bars denote the standard errors of the means. **C.** Pearson Correlation of the slopes for both sets. Each dot represents the slope for both sets from the same session. The slopes were transformed to the standard score before applying the correlation analysis. 95% confidence interval.

**Figure S3.**
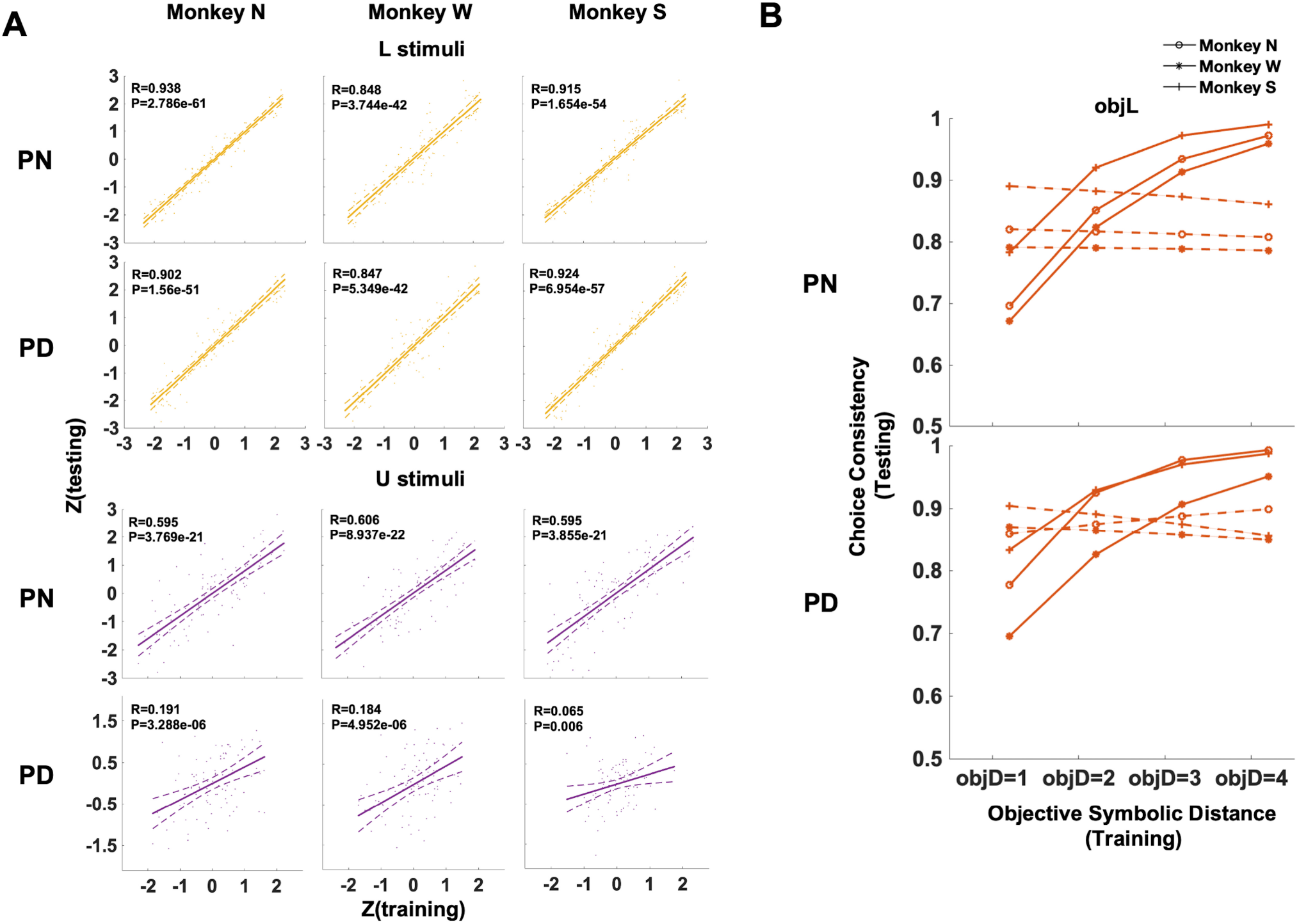
Subject showed consistent subjective ordering between training and testing. **A.** Pearson Correlation between the z-scores inferred from training and testing stage on each L(top) and U stimulus(bottom) under both PN and PD schedules, 95% confidence interval. **B.** Under both schedules, all subjects showed significant expected objective symbolic distance effect that disappeared when the objective distance was shuffled, indicating that subjects succeeded in inferring the order from the L set during training and applied it during testing persistently. Error bars denote the standard errors over all the posterior Bayesian regression coefficients. (top, PN, 95% CI for the objSD regressor N: [0.9055, 0.9105], W: [0.7907, 0.7954], S: [1.2809, 1.2889]; objSD regressor shuffled N: [-0.0509, −0.0473], W: [0.0039, 0.0076], S: [0.0482, 0.0529]; bottom, PD, 95% CI for the objSD regressor N: [1.0300, 1.0367], W: [0.7756, 0.7809], S: [1.4873, 1.4988]; objSD regressor shuffled N: [0.0681, 0.0726], W: [-0.005, −0.00066], S: [-0.0064, −0.0019]).

**Figure S4.**
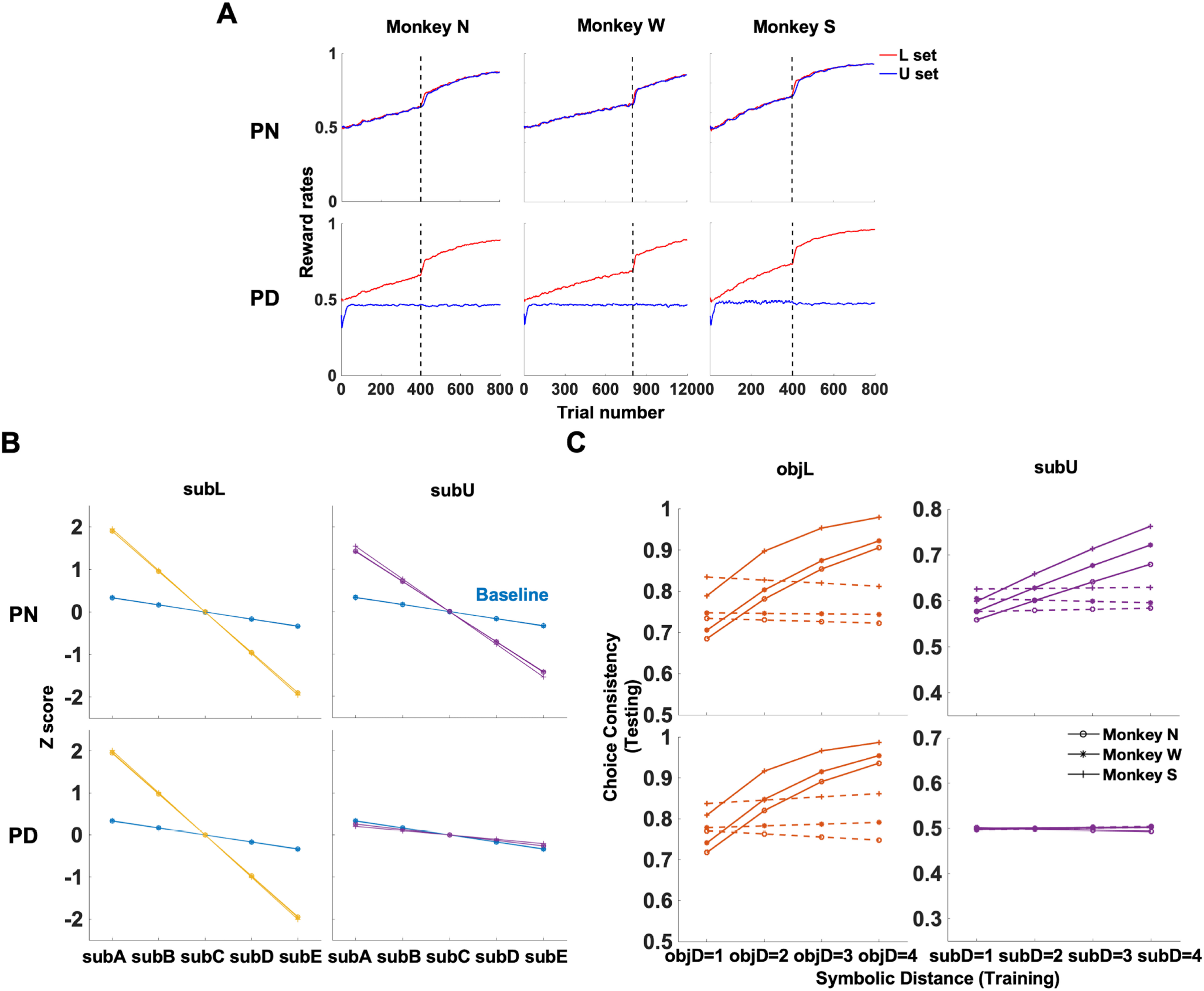
Model simulations. **A.** 10 L/U trials rolling reward rates over the course of the session for *Q*-learning under both schedules. Error bars denote the standard error of the means. The dash line demarcates the transition from training to testing stage **B.** The predicted ordering gradients from *Q*-learning for individual subject. SE (mean slope. Baseline: N: −0.1676, W: −0.1695, S: −0.1645; PN: L set, N: −0.9503, W: −0.9517, S: −0.9777; U set, N: −0.7154, W: −0.7083, S: −0.7704; PN: L set, N: −0.9719, W: −0.9804, S: −1.0038; U set, N: −0.0072, W: −0.1295, S: −0.1030) C. Predicted transfer effect from *Q*-learning for individual subject. Error bars referred to the standard errors over all the posterior Bayesian regression coefficients. The solid line refers to the original result while the dash line corresponds to the data with the distance shuffled across trials. (PN: mean L objSD regressor N: 0.8686, W: 0.774, S: 1.4067; objSD regressor shuffled N: −0.0072, W: 0.0324, S: 0.0024; U subSD regressor N: 0.178, W: 0.2327, S: 0.2576; subSD regressor shuffled N: 0.0002, W: 0.0055, S: 0.011; PD: L objSD regressor N: 0.9897, W: 0.9682, S: 1.7856; objSD regressor shuffled N: −0.035, W: 0.0159, S: −0.0061; U the subSD regressor N: −0.0055, W: 0.0068, S: −0.0002; subSD regressor shuffled N: 0.004, W: 0.0026, S: 0.0017)

**Figure S5.**
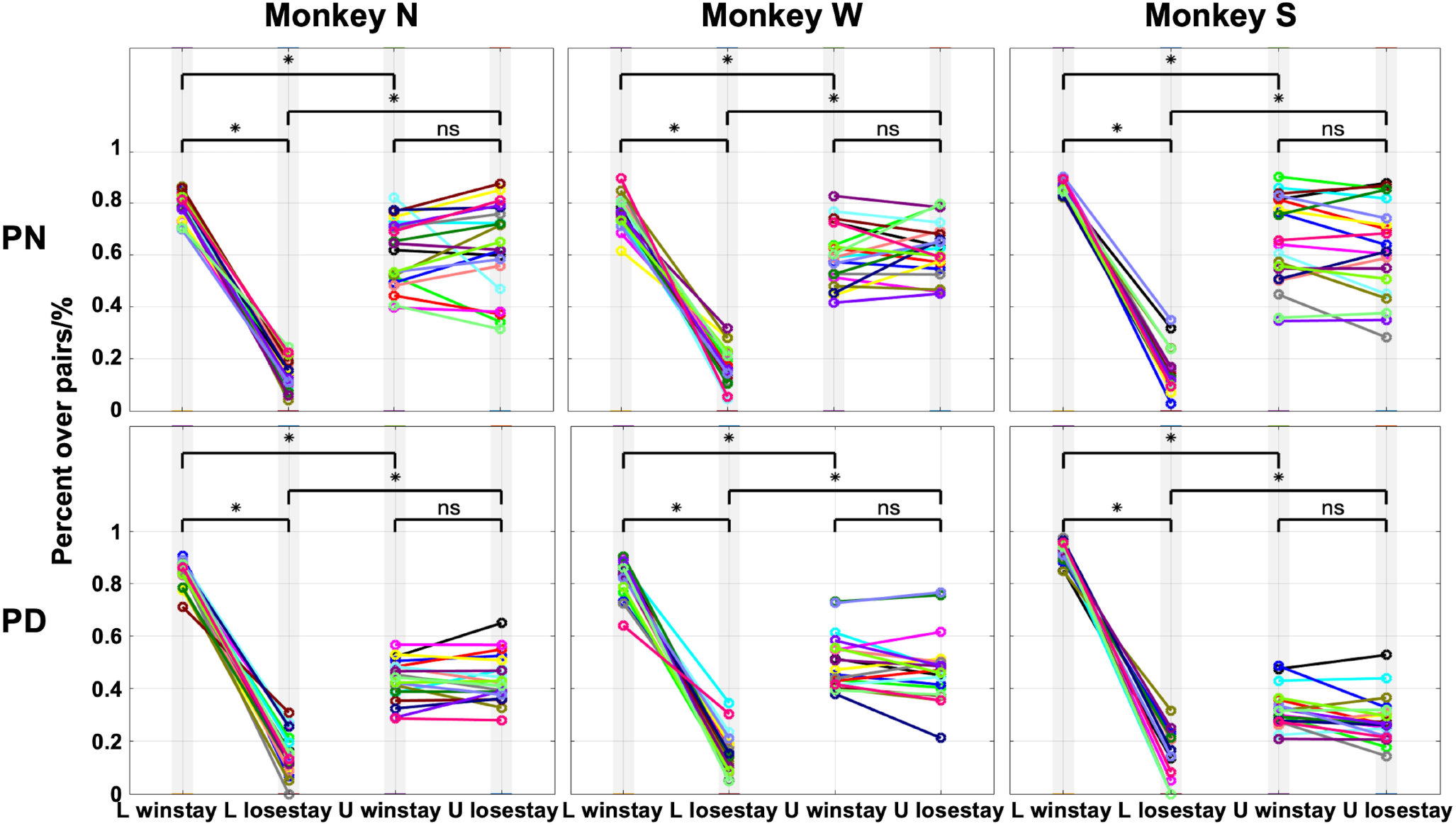
Monkeys exhibited different past-trial effects on L and U pairs. Under both schedules, all three subjects showed significantly higher percentage of win-stay than lose-stay over the L pairs whereas both strategies were indistinctively adopted over the U pairs during the testing stage. Each color corresponds to the data from one session. Shaded bar denotes each column of the data is significantly different from chance. *p* < .05, Wilcoxon signed rank test.

